# Whole-genome sequence and methylome profiling of the almond (*Prunus dulcis* [Mill.] D.A.Webb) cultivar ‘Nonpareil’

**DOI:** 10.1101/2021.10.27.466198

**Authors:** Katherine M. D’Amico-Willman, Wilberforce Z. Ouma, Tea Meulia, Gina M. Sideli, Thomas M. Gradziel, Jonathan Fresnedo-Ramírez

## Abstract

Almond (*Prunus dulcis* [Mill.] D.A. Webb) is an economically important, specialty nut crop grown almost exclusively in the United States. Breeding and improvement efforts worldwide have led to the development of key, productive cultivars, including ‘Nonpareil,’ which is the most widely grown almond cultivar. Thus far, genomic resources for this species have been limited, and a whole-genome assembly for ‘Nonpareil’ is not currently available despite its economic importance and use in almond breeding worldwide. We generated a 615.89X coverage genome sequence using Illumina, PacBio, and optical mapping technologies. Gene prediction revealed 27,487 genes using MinION Oxford nanopore and Illumina RNA sequencing, and genome annotation found that 68% of predicted models are associated with at least one biological function. Further, epigenetic signatures of almond, namely DNA cytosine methylation, have been implicated in a variety of phenotypes including self-compatibility, bud dormancy, and development of non-infectious bud failure. In addition to the genome sequence and annotation, this report also provides the complete methylome of several key almond tissues, including leaf, flower, endocarp, mesocarp, fruit skin, and seed coat. Comparisons between methylation profiles in these tissues revealed differences in genome-wide weighted percent methylation and chromosome-level methylation enrichment. The raw sequencing data are available on NCBI Sequence Read Archive, and the complete genome sequence and annotation files are available on NCBI Genbank. All data can be used without restriction.

## Introduction

In the past decade, significant advances have been made in the ability to produce whole-genome sequences from *Prunus* species. Challenges due to the inherent characteristics of outcrossing heterozygous plants have prevented the *Prunus* community from producing high-quality assemblies using second-generation high-throughput sequencing (Illumina) and pyrosequencing (Roche 544) technologies. As a result, the first *Prunus* genome was produced using a doubled-haploid of the peach rootstock genotype ‘Lovell’ (The International Peach Genome Initiative *et al*. 2013). Although this ‘Lovell’ genome was further refined using short-read sequencing data to ensure continuity and completeness (Verde *et al*. 2017), the capabilities for developing genome assemblies for heterozygous *Prunus* genotypes were very limited. These limitations excluded other commercially relevant cultivars from being sequenced at that time.

The recent advent, improvement, and successful implementation of third-generation high-throughput sequencing technologies such as single-molecule real-time sequencing (Pacific Biosciences) and Nanopore Sequencing (Oxford) has been key to sequencing heterozygous *Prunus* genomes. At the time of preparing this manuscript, twenty genomes assemblies for thirteen *Prunus* species (https://www.rosaceae.org/species/prunus/all) are reported in the Genome Rosaceae Database (GDR, www.rosaceae.org) (Jung *et al*. 2019). Among these new resources are two *P. dulcis* genomes: one from the cultivar ‘Lauranne’ (Sánchez-Pérez *et al*. 2019), which was used to investigate kernel bitterness, and the other from the cultivar ‘Texas’ (a.k.a. ‘Mission’) (Alioto *et al*. 2019), which was used to study the nature of transposable elements and their contribution to structural divergence from peach.

Since 2018, our group has worked to produce a genome assembly for ‘Nonpareil,’ the most widely grown almond genotype. ‘Nonpareil’ was first described in 1879 and currently represents 42% of US almond production along with significant use in other almond-producing countries such as Australia (Almond Board of California 2020). ‘Nonpareil’ is considered the reference cultivar in terms of field performance and kernel quality for a large portion of the industry and is therefore used as a recurrent parent in almond breeding programs. However, ‘Nonpareil’ is susceptible to the aging-related disorder noninfectious bud-failure (BF), which can negatively impact kernel yield and has been associated with genome-wide DNA methylation (Fresnedo-Ramírez *et al*. 2017). Our group developed a genome assembly as well as methylomes for a variety of almond tissues using this relevant cultivar to address BF by assessing the risk of onset in almond breeding material. Developing and exploring *Prunus* genomes such as ‘Nonpareil’ will shed light on the genome polymorphism and dynamism involved in the exhibition of agriculturally relevant traits and syndromes.

## Methods & Materials

### Plant material

Tissue samples for the genome sequencing and methylome profiling were collected in 2018, 2019, and 2020 from a single ‘Nonpareil’ clone: GOH B32 T37-40 maintained at Foundation Plant Services – University of California, Davis (Davis, CA, USA). This ‘Nonpareil’ clone is recognized as the foundational source for any ‘Nonpareil’ almond deployed in commercial orchards worldwide. Leaf samples were collected in 2018 for whole-genome sequencing and for RNA sequencing from another ‘Nonpareil’ clone located at the Wolfskill Experimental Orchards – University of California, Davis (Winters, CA, USA) which is grafted onto ‘Nemaguard’ peach rootstock and exhibits non-infectious bud failure.

Initially, leaf samples were collected for whole-genome and RNA sequencing in 2018. In 2019, additional leaf samples were collected for optical mapping, while phloem tissue and fruits were collected for DNA methylation profiling and RNA sequencing. In 2020, flower tissues were collected for RNA sequencing and DNA methylation profiling prior to anthesis. Leaf tissues were collected on ice in the field and stored at −20° C until sample processing, while flower tissue was collected and immediately put on dry ice and stored at −80° C until processing. Leaf samples collected in 2018 and 2019 were processed at UC Davis, and additional samples were shipped overnight on dry ice in 2018 to the Ohio Agricultural Research and Development Center (OARDC – Wooster, Ohio) and stored at −40° C until processing for RNA sequencing. Fruits and phloem tissues collected in 2019 were shipped overnight on ice to the OARDC and stored at −20° C until sample processing. Flower tissues were processed at UC Davis for RNA sequencing, and additional samples were shipped overnight on ice to the OARDC for DNA methylation profiling.

### DNA isolation

For long-read and short-read sequencing, leaf samples were sent to Dovetail Genomics®, where DNA was isolated using a standard CTAB protocol. For optical mapping, ultra-high molecular weight (uHMW) DNA was isolated from leaves using the Plant DNA Isolation Kit (Bionano Genomics, San Diego, CA) according to the manufacturer’s instructions. The uHMW DNA molecules were labeled with the DLE-1 enzyme (Bionano Genomics, San Diego, CA) and stained using the Bionano Prep™ Direct Label and Stain (DLS) Kit (Bionano Genomics, San Diego, CA) according to the manufacturer’s instructions.

For methylation profiling, DNA was isolated from young leaves, fruits, and flowers. To extract DNA from fruit, the fruits were first dissected using a scalpel while frozen to isolate fruit skin, mesocarp, endocarp, and seed coat tissues. All tissues were then ground in liquid nitrogen using mortar and pestle, and 150 mg of ground material was used as input into the SILEX DNA isolation protocol outlined in Vilanova *et al*. (2020) with some modifications. To isolate DNA from young leaves, leaf tissue was ground to a fine powder in liquid nitrogen using a mortar and pestle, and 150 mg of ground material was used as input into the SILEX DNA isolation protocol as above. Finally, DNA was isolated from whole flowers using the same method of grinding in liquid nitrogen and isolating following the SILEX protocol. All isolated DNA was assessed for concentration and quality by fluorometry and electrophoresis using a Qubit™ 4 and Qubit™ 1X dsDNA HS Assay Kit (ThermoFisher Scientific) and a TapeStation (Agilent, Santa Clara, CA, USA).

### RNA isolation, library preparation, and sequencing

To isolate RNA from leaf, fruit, and phloem tissues samples, tissue was ground in liquid nitrogen using a mortar and pestle, and 50 mg of tissue was used as input into the RNA isolation protocol outlined in Gambino *et al*. (2008)(Gambino *et al*. 2008). To isolate RNA from flowers, the tissues were ground in liquid nitrogen, and the ground material was used as input in ThermoFisher Scientific PureLink™ Plant RNA Reagent. Following extraction, RNA was DNase treated using the DNA-*free*™ DNA Removal Kit (ThermoFisher Scientific) according to the manufacturer’s instructions. RNA concentration and quality were assessed by fluorometry and electrophoresis using a Qubit™ 4 and Qubit RNA HS Assay Kit (ThermoFisher Scientific) and a TapeStation (Agilent).

A sequencing library for the flower RNA sample was prepared using the NEBNext^®^ Ultra™ II Directional RNA Library Prep Kit for Illumina^®^ (New England BioLabs^®^, Inc., Ipswich, MA, USA) and barcoded using index primers from the NEBNext^®^ Multiplex Oligos for Illumina^®^ (New England BioLabs^®^) following the manufacturer’s instructions. The library was equimolarly pooled and split for sequencing on two lanes of the Illumina^®^ HiSeq 4000 in paired-end 2 × 150-bp mode.

The RNA-seq libraries for short-read sequencing for the fruit, leaf, and phloem tissues were also prepared using the NEBNext^®^ Ultra^™^ II Directional RNA Library Prep Kit for Illumina^®^ (New England BioLabs^®^) and barcoded using index primers from the NEBNext^®^ Multiplex Oligos for Illumina^®^ (New England BioLabs^®^). These libraries were processed using the MiSeq Reagent Kit v2 according to the manufacturer’s protocols and were sequenced in an Illumina MiSeq device in paired-end 2 × 150-bp mode.

### Sequencing of RNA-seq libraries using Oxford Nanopore technology (ONT)

Total RNA from leaves was first depleted using the Illumina^®^ Ribo-Zero rRNA Removal Kit (Plant). The RNA was then purified and concentrated using the RNA Clean Concentrator™-5 kit (Zymo Research, Irvine, CA, USA). cDNA libraries were prepared using a mix of 50 ng RNA and 0.5 ng Spike-in RNA Variant Control Mix E2 (Lexogen, NH, USA) according to the Oxford Nanopore Technologies (Oxford Nanopore Technologies Ltd, Oxford, UK) protocol “DNA-PCR Sequencing” with 14 cycles of PCR (8-minute elongation time). ONT adapters were ligated to 650 ng of cDNA. These libraries were sequenced using a MinION Mk1b with an R9.4.1 flowcell. The data were preprocessed using MinKNOW 3.1.18, and the base calling was done using Guppy 2.0.5.

### Library construction, sequencing, and assembly of PacBio data

Long-read sequencing was performed with a PacBio Sequel II System (Pacific Biosciences, Menlo Park, CA, USA) using Single Molecule, Real-Time (SMRT) technology. Genomic DNA libraries were prepared with 5 μg of input DNA according to the “Guidelines for Preparing 20 kb SMRTbell™ Templates” (available at https://www.pacb.com/wp-content/uploads/2015/09/User-Bulletin-Guidelines-for-Preparing-20-kb-SMRTbell-Templates.pdf). Sequencing was performed on one PacBio Sequel II 8M SMRT cell by Dovetail Genomics^®^ (Santa Cruz, CA, USA). The data yielded from the long-read sequencing was processed for *de novo* assembly using the FALCON v. 0.3.0 (Chin *et al*. 2016) pipeline customized by Dovetail Genomics^®^ for the assembly of a heterozygous genome. The assembly was polished using Arrow v. 2.3.3 (available at https://github.com/PacificBiosciences/pbbioconda).

### Optical map construction

A consensus optical map was assembled *de novo* with the assembler tool in the Bionano Solve v. 3.4 package (available at https://bionanogenomics.com/support/software-downloads/) using significance cutoffs of P < 1 × 10^−8^ to generate draft consensus contigs, P < 1 × 10^−9^ for draft consensus contig extension, and P < 1 × 10^−15^ for the final merging of the draft consensus contigs; a recipe of “haplotype,” “noES,” and “noCut” was chosen for the assembly. The initial optical map was then checked for potential chimeric contigs and further refined.

### Scaffolding

The sequence assembly was validated by comparing it with the optical map. The contigs of the PacBio assembly were digested *in silico* with DLE-1 restriction sites using Knickers (Bionano Genomics, San Diego, CA, USA). The alignment was performed using the RefAligner tool in the Bionano Solve v 3.4 package (Bionano Genomics, San Diego, CA) with an initial alignment cutoff of P < 1 × 10^−10^. The PacBio contigs that disagreed with the optical map were disjoined accordingly, and the conflict-free contigs were linked with the guide of the optical map. The gaps were then filled with the respective number of N’s using the estimated length between the flanking restriction sites.

### Pseudomolecule construction

Genotyping-by-sequencing data generated for a mapping population of 89 individuals (Goonetilleke *et al*. 2018) were previously used to produce a linkage map using the software package RABBIT v. 3.2. (Zheng *et al*. 2019) (https://github.com/chaozhi/RABBIT). The linkage map was compared with the assembly and gene annotations of *Prunus dulcis* cv. ‘Texas’ v. 2.0 (Alioto *et al*. 2019) to confirm genome collinearity. The linkage map was then used to determine the order and orientations of the scaffolds on each chromosome. The scaffolds were linked with 100 N’s accordingly and anchored onto the eight chromosomes.

### Gene prediction and annotation

Repeats and low complexity regions were identified and subsequently masked in the ‘Nonpareil’ assembly. Illumina reads generated from the short-read RNA sequencing of both the flower and leaf tissue were assembled with Trinity software (Grabherr *et al*. 2011; Haas *et al*. 2013) prior to training the SNAP gene finder classifier (Korf 2004). An iterative training process resulted in the generation of a SNAP hidden Markov model (HMM), which was subsequently employed—together with an AUGUSTUS gene finder—in predicting an initial list of gene models (Haas *et al*. 2013). Long-reads from cDNA sequenced using ONT were first trimmed using porechop v.0.2.4 (https://github.com/rrwick/Porechop) and then aligned to the newly assembled transcriptome generated from the Illumina reads using Minimap2 v. 3.16 (Li 2018).

For functional annotation, predicted gene models were submitted to a pipeline utilizing database searches of protein sequences in Uniprot (homology search) (Consortium 2019), KEGG database (orthology search) (Kanehisa and Goto 2000), and Pfam (Consortium 2019). The resulting descriptions of putative gene functions, Pfam domain identifiers (IDs), and gene ontology (GO) terms were included in the genome annotation feature file (GFF).

### Enzymatic methyl-seq library preparation and sequencing

Whole-genome enzymatic methyl-seq libraries were prepared using the NEBNext® Enzymatic Methyl-seq kit (New England BioLabs®, Inc.) following the protocol for standard insert libraries (370-420 base pairs). Each sample was prepared using 100 ng input DNA in 48 μL TE buffer (1 mM Tris-HCl; 0.1 mM EDTA; pH 8.0) with 1 μL spikes of both the CpG unmethylated Lambda and CpG methylated pUC19 control DNA provided in the kit. The samples were sonicated using a Covaris® S220 focused-ultrasonicator in microTUBE AFA Fiber Pre-Slit Snap-Cap 6×16 mm tubes (Covaris®, Woburn, MA, USA) with the following program parameters: peak incident power (W) = 140; duty factor = 10%; cycles per burst = 200; treatment time (s) = 80.

Following library preparation, library concentration and quality were assessed by fluorometry using a Qubit™ 4 and Qubit™ 1X dsDNA HS Assay Kit (ThermoFisher Scientific) and by electrophoresis using a TapeStation (Agilent). Library concentration was further quantified by qPCR using the NEBNext® Library Quant Kit for Illumina® (New England BioLabs®, Inc.). Libraries were sequenced on one lane of the Illumina® HiSeq4000 platform to generate 150-bp paired-end reads.

### Processing and alignment of enzymatic methyl-seq libraries

Methyl-Seq read quality was initially assessed using FastQC v. 0.11.7 (Andrews 2010), and reads were trimmed using TrimGalore v. 0.6.6 and Cutadapt v. 2.10 with default parameters (Krueger 2016). Forward read fastq and reverse read fastq files from the two HiSeq4000 lanes were combined for each library to produce single fastq files for both read one and read two. Reads were aligned to the ‘Nonpareil’ v. 2.0 almond reference genome, deduplicated, and methylation calls were generated using Bismark v. 0.22.3 (Krueger and Andrews 2011) with default parameters in paired-end mode. Reads were also aligned to both the Lambda and pUC19 nucleotide sequence fasta files provided by NEB (https://www.neb.com/tools-and-resources/interactive-tools/dna-sequences-and-maps-tool) to test conversion efficiency. All analyses were performed using the Ohio Supercomputer Center computing resources (Ohio Supercomputer Center, 1987).

### Analysis of DNA methylation profiles in almond tissues

Weighted genome-wide percent methylation values were calculated for each tissue type by taking the total number of methylated reads at each cytosine and dividing this by the total number of reads (methylated + unmethylated) at each cytosine. Weighted values were calculated for each methylation context. Methylation call files were then subset by chromosome (chr1 – chr8), and weighted percent methylation values were calculated for each tissue type by chromosome using the same formula as above for each methylation context. To visualize weighted percent methylation across the eight chromosomes in the ‘Nonpareil’ genome, circos plots were generated with one track depicting the weighted percent methylation for each of five tissue types (leaf, fruit skin, mesocarp, endocarp, and seedcoat) across the genome. To create the circos plots, the R package circlize v. 0.41.2 (Gu *et al*. 2014) was used along with text files containing aggregated percent methylation values generated using bedtools *makewindows* with a bin size of 100,000 base pairs (Quinlan and Hall 2010). The command *circos.genomicTrack()* was used to create each track depicting weighted percent methylation for each tissue type across the eight chromosomes of the almond genome (Gu *et al*. 2014).

### Data Availability

Raw sequencing read data files for the RNA sequencing, enzymatic methyl-seq, short-read whole-genome sequencing, and long-read whole-genome sequencing are all available through the NCBI Sequence Read Archive under the Bioproject <u>PRJNA769745</u>. The genome assembly is available in the NCBI GenBank under the accession number JAJFAZ000000000. Supplementary files for the optical mapping are also available at NCBI under the SUPPF_0000004114.

## Results and Discussion

The first version of the almond ‘Nonpareil’ genome assembly (available at: https://www.rosaceae.org/rosaceae_downloads/Prunus_dulcis/Nonpareil_v1.tar.gz) was the result of a combination of Illumina technology (HiSeq X) and Hi-C data in the form of CHiCAGO and Dovetail HiRise® implementations. With a coverage of 3739.9X (assuming a genome size of 240 Mb), the resulting assembly of 164.55 Mb represented the gene space of ‘Nonpareil’ assembled in 2081 scaffolds with N50 =15.28 Kb. Using Hi-C, it was possible to infer the eight major pseudomolecules expected in the haploid assembly representing the genome space of ‘Nonpareil.’

Here, we present the results of a second iteration of the assembly and genome annotation of ‘Nonpareil.’ The purpose was to improve the representation, completeness, and orientation of the genome using long-read sequencing coupled with optical mapping technology. Approximately 793 million reads were produced from PacBio CLR libraries sequenced on the PacBio Sequel II, yielding approximately 147 Gb of data with a mean read length of 23.3 kb. These reads represent a coverage of 615.89X (assuming a genome size of 240 Mb). A total of 784.27 million corrected reads were used as input in the PacBio FALCON whole-genome assembly pipeline. The N50 length of the error corrected reads was 29.33 kb. The final assembly had a contig N50 of 1.34 Mb and N90 of 384.64 kb. The final number of polished contigs was 593 encompassing 456.66 Mb with a 97.1% BUSCO completeness score and an NG50 of 2.83 Mb.

This assembly was used as input for optical mapping for which approximately 52 Gb of large single molecules (> 150 Kb), labeled by DLE-1 enzyme (Bionano Genomics), were collected and *de novo* assembled into an optical map. This map consists of 227 contigs with a total length of 533.72 Mb and an N50 of 3.19 Mb. We aligned the 456.66 Mb PacBio contigs (Contig_v1.0; Table 1) of almond to the optical map and identified 244 conflicts included in 194 contigs (totaled 288,407,142 bp). These chimeric contigs were resolved, and the redundancies were removed. The conflict-free dataset (Contig_v1.1) consists of 740 contigs with a slightly decreased N50 size of 1,308,716bp (Table 1). The sequences of Contig_v1.1 were used to generate scaffolds with the optical map as a guide resulting in scaffolds with a total length of 458,275,742 bp and N50 size of 3,168,198 bp (Scaffold_all; Table 1). The longest scaffold size is 27,216,209 bp.

**Table 1.**
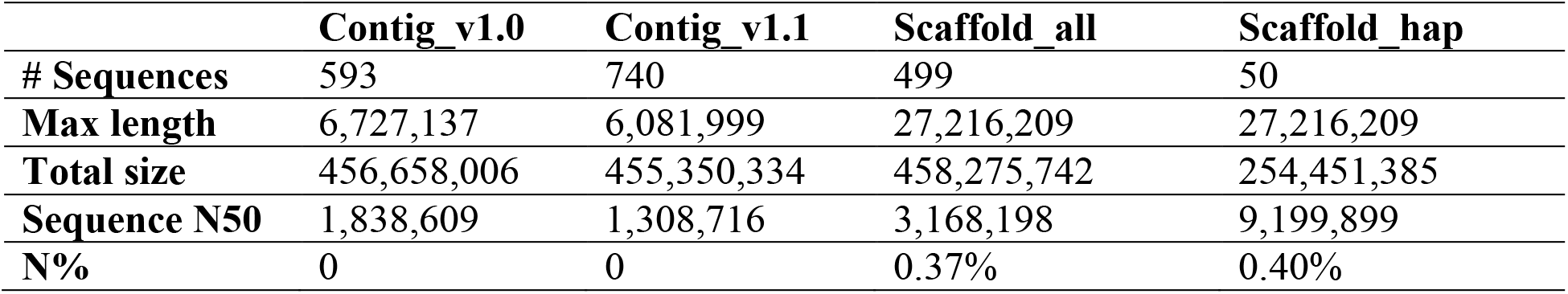
Summary of scaffolding using the optical map.

|  | **Contig_v1.0** | **Contig_v1.1** | **Scaffold_all** | **Scaffold_hap** |
| --- | --- | --- | --- | --- |
| **# Sequences** | 593 | 740 | 499 | 50 |
| **Max length** | 6,727,137 | 6,081,999 | 27,216,209 | 27,216,209 |
| **Total size** | 456,658,006 | 455,350,334 | 458,275,742 | 254,451,385 |
| **Sequence N50** | 1,838,609 | 1,308,716 | 3,168,198 | 9,199,899 |
| **N%** | 0 | 0 | 0.37% | 0.40% |

The optical map was self-aligned to obtain two datasets, each representing a haplotype. As a result, two datasets with total sizes of approximately 275 Mb and 258 Mb, respectively, were generated. The Scaffold_all was divided into two datasets accordingly by aligning sequences with the two sets of optical maps. The primary scaffold set (Scaffold_hap; Table 1) has a total size of 254,451,385 bp and an N50 of 9,199,899 and was used to construct pseudomolecules.

We then used the linkage map as well as the gene annotations of the *Prunus dulcis* cv. ‘Texas’ v. 2.0 to order and orient these scaffolds containing two or more segregating linkage markers or genes. We managed to anchor all 50 scaffolds of Scaffold_hap onto the eight chromosomes. The characteristics of each pseudomolecule are summarized in Table 2. This genome assembly was processed for deposit in GenBank by trimming all possible organellar contaminations in the nuclear genome. The functional annotation (as a GFF file) was also deposited as additional data.

**Table 2.** Summary of the pseudomolecules.

| **Chromosome** | **Length (bp)** | **Effective length (bp)** | **N%** |
| --- | --- | --- | --- |
| **Pdu1** | 53,051,038 | 52,635,155 | 0.78 |
| **Pdu2** | 33,983,478 | 33,651,924 | 0.98 |
| **Pdu3** | 30,551,305 | 30,359,259 | 0.63 |
| **Pdu4** | 30,646,815 | 30,645,741 | 0.00 |
| **Pdu5** | 21,034,137 | 21,021,108 | 0.06 |
| **Pdu6** | 32,795,989 | 32,735,446 | 0.18 |
| **Pdu7** | 26,339,959 | 26,327,521 | 0.05 |
| **Pdu8** | 26,052,964 | 26,049,868 | 0.01 |
| **Total** | 254,455,685 | 253,426,022 | 0.40 |

Using the genome generated in this report, we analyzed whole-genome DNA methylation profile data for six tissue types (leaf, flower, fruit skin, mesocarp, endocarp, and seed coat) from ‘Nonpareil.’ Based on this analysis, we found genome-wide weighted total percent methylation values for each methylation context (CG, CHG, CHH [H = C, T, or A]) (Table 3). Total percent weighted methylation is lowest in the CHH context, as has been reported in other angiosperms (Niederhuth *et al*. 2016). Some variation in genome-wide methylation values was observed across tissue types, with leaf tissue exhibiting the lowest overall methylation levels of the tissues tested. Interestingly, flower tissue showed the highest methylation levels in the CG and CHG context but was observed to have the second lowest methylation level in the CHH context (Table 3).

**Table 3.** Genome-wide weighted percent methylation values for each tissue type in ‘Nonpareil.’ Weighted percent methylation is depicted for each methylation context (CG, CHG, and CHH [H = A, T, or C]).

| **Tissue Type** | **% CG** | **% CHG** | **% CHH** |
| --- | --- | --- | --- |
| **Leaf** | 45.2 | 14.6 | 1.3 |
| **Flower** | 56.4 | 29.4 | 2.9 |
| **Fruit Skin** | 52.5 | 17.7 | 4.0 |
| **Mesocarp** | 52.9 | 18.3 | 4.7 |
| **Endocarp** | 51.7 | 17.7 | 4.3 |
| **Seed Coat** | 50.5 | 16.7 | 4.3 |

Genome-wide methylation was also displayed across the eight chromosomes of the almond genome in each of the three methylation contexts (**CG** – Fig. 1a; **CHG** – Fig. 1b; **CHH** – Fig. 1c). The circos plots show similar patterns in the distribution of methylation across the almond genome for each tissue type tested (Fig. 1a-c). Variations in methylation profiles across tissue types in plants have been previously found to be low, particularly in non-embryonic tissues (Schmitz *et al*. 2013; Kawakatsu *et al*. 2016)(Schmitz *et al*., 2013; Kawakatsu *et al*., 2016).

**Figure 1.**
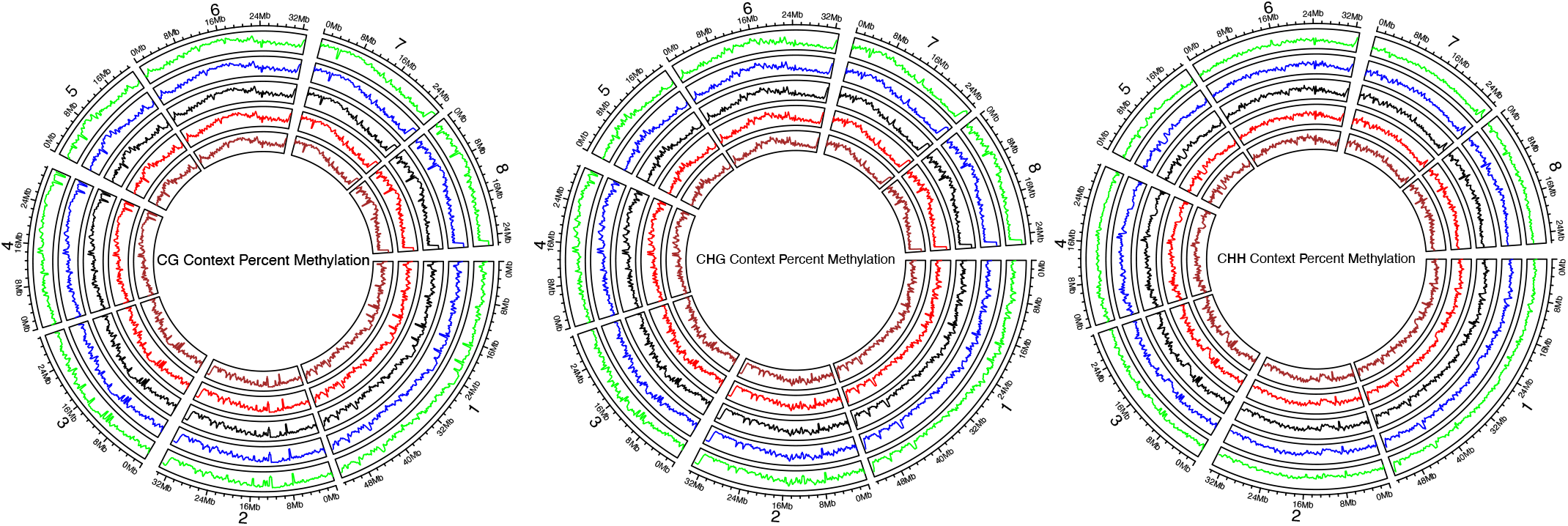
Circos plots depicting genome-wide weighted methylation across each of the 8 almond chromosomes for each methylation context (CG, CHG, and CHH [H = A, T, or C]). Five almond tissue types are represented in each genomic track: Leaf (green), Fruit Skin (blue), Mesocarp (black), Endocarp (red), and Seedcoat (brown).

The ‘Nonpareil’ genome sequence and whole genome methylome data generated in this report represent a continued commitment of the *Prunus* community to improve the availability of genetic resources for these economically valuable species. Further, this report presents the first genome assembly for ‘Nonpareil,’ the most widely grown almond cultivar in the US. Access to these data provides researchers in public and private institutions the ability to use genomics to improve and protect almond production, as well as address future challenges imposed by climate change, newly introduced pests or pathogens, and the volatility of consumer preference.

## Acknowledgements

The Ohio State University CFAES-SEEDS program grant 2019-125, the Almond Board of California Grant HORT35, the Translational Plant Sciences Graduate Fellowship, the Cancer Center Support Grant (CCSG) P30CA016058, the AFRI-EWD Predoctoral Fellowship 2019-67011-29558 from the USDA National Institute of Food and Agriculture, and the Ohio Supercomputer Center. We would like to thank Elizabeth S. Anderson for her help in the preparation of samples and extraction of RNA from distinct fruit tissues. Also, thanks to Dr. Shashi N, Goonetilleke, and Prof. Diane E. Mather at The University of Adelaide for providing access to the keyfile for the GBS data generated from the ‘Nonpareil’ × ‘Lauranne’ mapping population.

